# Metacommunities from bacteria to birds: stronger environmental selection in mediterranean than in tropical ponds

**DOI:** 10.1101/2021.03.24.436596

**Authors:** Ángel Gálvez, Pedro R. Peres-Neto, Andreu Castillo-Escrivà, Fabián Bonilla, Antonio Camacho, Eduardo M. García-Roger, Sanda Iepure, Javier Miralles, Juan S. Monrós, Carla Olmo, Antonio Picazo, Carmen Rojo, Juan Rueda, María Sahuquillo, Mahmood Sasa, Mati Segura, Xavier Armengol, Francesc Mesquita-Joanes

**Affiliations:** Cavanilles Institute for Biodiversity and Evolutionary Biology, University of Valencia, Paterna, Valencia, Spain; Department of Biology, Concordia University, Montreal, Canada; Instituto Clodomiro Picado, Facultad de Microbiología, Universidad de Costa Rica, San José, Costa Rica; Emil Racovitza Institute of Speleology, Cluj Napoca, Romania; Subdirecció General del Medi Natural, Generalitat Valenciana, València, Spain; Museo de Zoología, Centro de Investigación en Biodiversidad y Ecología Tropical, Universidad de Costa Rica, Costa Rica

**Keywords:** multi-taxon analysis, dispersal limitation, environmental selection, tropical and temperate ecology, freshwater metacommunity

## Abstract

The metacommunity concept provides a theoretical framework that aims at explaining organism distributions by a combination of environmental filtering, dispersal and drift. With the development of statistical tools to quantify and partially isolate the role of each of these processes, empirical metacommunity studies have multiplied worldwide. However, few works attempt a multi-taxon approach and even fewer compare two distant biogeographical regions using the same methodology. Under this framework, we tested the expectation that temperate (mediterranean-climate) pond metacommunities would be more influenced by environmental and spatial processes than tropical ones, because of stronger environmental gradients and greater isolation of waterbodies.

We surveyed 30 tropical and 32 mediterranean temporary ponds from Costa Rica and Spain, respectively, and obtained data on 49 environmental variables (including limnological, hydrogeomorphological, biotic, climatic, and landscape variables). We characterized the biological communities of Bacteria and Archaea (from both the water column and the sediments), phytoplankton, zooplankton, benthic invertebrates, amphibians and birds, and estimated the relative role of space and environment on metacommunity organization for each group and region, by means of variation partitioning using Generalized Additive Models (GAMs).

Environmental selection was important in both tropical and mediterranean ponds, but markedly stronger in the latter, probably due to their larger limnological heterogeneity. Spatialized environment and pure spatial effects were greater in the tropics, related to higher climatic heterogeneity and dispersal processes (e.g. restriction, surplus) acting at different scales. The variability between taxonomic groups in spatial and environmental contributions was very wide. Effects on passive and active dispersers were similar within regions but different across regions, with higher environmental effects in mediterranean active dispersers. The residual (unexplained) variation was larger in tropical pond metacommunities, suggesting a higher role for stochastic processes and/or effects of biotic interactions in the tropics. Overall, these results provide support, for a wide variety of organisms related to aquatic habitats, for the classical view of stronger abiotic niche constraints in temperate areas compared to the tropics.

---

> “Tropical species often exhibit patchy geographic distributions that respond to no obvious relations with climate or habitat”
>
> — MacArthur, 1972

## Introduction

Ecological communities are not isolated systems, as they were often considered in the past, but are instead components of large spatial networks connected by dispersal, known as metacommunities (Hanski and Gilpin 1991; Wilson 1992). Although metacommunity assembly involves intricate mechanisms and processes, these can be classified into three broad sets, involving environmental selection, dispersal and stochastic processes (Vellend 2010; Leibold and Chase 2018). Extreme scenarios of these processes are often conceptualized under four archetypes, namely species sorting, patch dynamics, mass effects and neutral models (Leibold et al. 2004). Research in the past two decades has established that the processes underlying metacommunity structure depend on complex interactions between local patch attributes and landscape settings (e.g., connectivity, environmental heterogeneity among sites; Grönroos et al. 2013; Erös et al. 2017; Castillo-Escrivà et al. 2017a), and species attributes (e.g., body size, dispersal mode, trophic level; Vanschoenwinkel et al. 2010; De Bie et al. 2012; Astorga et al. 2012). Our ability to detect, differentiate and understand the roles of different processes have been particularly challenging because of the complex ways that landscape and species characteristics may affect local community assembly and metacommunity structure. Despite the complexity in which assembly processes may take place, these are often conceptualized at taking place across two broad scales (Peres-Neto et al. 2012): regional processes (including biogeography; Leibold et al. 2010) that regulate the movement of organisms among local communities (e.g., landscape heterogeneity, connectivity, dispersal limitation); and local processes that regulate the success of species following either their own arrival or the arrival of other species (e.g., niche differentiation, local environment, microhabitat heterogeneity).

Our understanding of metacommunity is, more often than not, scale dependent (e.g., stream, basin and ecoregion; Heino et al. 2015; Leibold and Chase 2018). For instance, small spatial scales, with dispersal surplus and/or homogeneous environmental conditions, favor metacommunity homogeneity, whereas at large spatial scales, dispersal rates decrease, generating metacommunity variation consistent with dispersal limitation and wider environmental gradients. Both small and large spatial scales, however, may reduce the strength of environmental selection and/or environmental tracking, which should be more relevant at intermediate scales. This is because intermediate scales should be large enough to encompass heterogenous environments, yet allowing moderate dispersal rates to track suitable localities. As such, keeping spatial extent constant is somewhat essential for contrasting metacommunity structure across different landscapes and/or taxonomic groups (as spatial effects also depend on the dispersal abilities of each organism; Wiens 1989). Few studies, however, have explored metacommunity structure contrasting distinct biogeographical regions (with large differences in their abiotic settings and biota) while controlling for spatial extent (Myers et al. 2013; Heino et al. 2017).

More uniform environmental conditions in tropical regions should lead to reduced species sorting compared to the more heterogeneous environment in temperate areas (Leibold and Chase 2018). Indeed, some studies found that temperate communities were more environmentally controlled than tropical ones (Myers et al. 2013; Souffreau et al. 2015). In the case of aquatic habitats, massive floods during the rainy season increase aquatic connectivity in tropical areas (Junk et al. 1989; Bunn et al. 2006; Larsen et al. 2019), reducing dispersal limitation and homogenizing local communities (Thomaz et al. 2007; Brasil et al. 2020), and contrasting with the more intense isolation of water bodies in the mediterranean-climate region. In both tropical and mediterranean systems, we may then expect that metacommunity structure is a result of dynamic switches between dispersal limitation and environmental filtering through time (Jacobson and Peres-Neto. 2010; Fernandes et al. 2014).

Metacommunity studies focusing on distinct organisms within the same landscape and localities revealed strong differences among taxa regarding dispersal limitation versus environmental selection (Beisner et al. 2006; Padial et al. 2014; Gálvez et al. 2020). Variation in dispersal mode and body size have been considered important features associated with dispersal capacity and, consequently, modulating the strength of environmental filtering and dispersal limitation (De Bie et al. 2012). For active dispersers, dispersal ability increases with body size, being easier for large organisms to spread among suitable sites (favoring species sorting), thus avoiding dispersal limitation (Grönroos et al. 2013; Csercsa et al. 2019). Passive transport, on the other hand, may limit dispersal capacity of large-sized organisms in contrast to small ones (which also produce more propagules; Finlay 2002). As a result, environmental selection should in principle be stronger for small organisms (Van der Gucht et al. 2007) though the evidence about this association is mixed (e.g., Heino et al. 2012; Schulz et al. 2012). Taken together, dispersal capacity, species attributes, geographic variation in environmental features and biogeographic features should interact in structuring metacommunities (see Leibold et al. 2010 for a discussion).

One way to address the complexity in which local communities assemble locally and metacommunity dynamics take place is by determining whether: 1) Different taxa (including completely different modes of life, e.g., bacteria versus birds) assemble differently within the same landscape; or 2) different environmental settings (landscapes) structure different taxa in similar ways. To answer these questions, one needs to study the metacommunity structure of multiple taxa across different landscapes. While this approach can provide strong inferences about the generality (or lack thereof) of assembly mechanisms and whether contingencies are due to landscapes or species attribute differences, there are obvious logistic challenges. In this study, we set out an ambitious empirical study to contrast the relative importance of environmental *versus* spatial factors on structuring metacommunities. We sampled temporary pond metacommunities in an extremely wide range of taxonomic groups (across 24 groups of organisms in total, varying from bacteria to vertebrates) across two very distinct biogeographic regions (Neotropical and Mediterranean) within the same temporal span and encompassing the same spatial extent within regions, based on the same sampling design, protocols and data measurements. In both regions, Costa Rica and Spain, precipitations are distributed seasonally, with a marked dry season that promotes the formation of temporary ponds. These are valuable ecosystems for metacommunity studies due to their well delimited spatial and temporal boundaries, and their isolation within a terrestrial matrix (Leibold and Chase 2018; at least for aquatic organisms). We framed our study around three predictions: 1) both environmental and spatial effects should have lower influences on metacommunity structure of tropical in contrast to mediterranean ponds; 2) different taxa should respond differently within regions but each similarly between regions, and in particular 3) metacommunity structure should be related to variation of dispersal abilities among taxa, in which higher environmental filtering should occur for small passive dispersers and large-bodied active dispersers in contrast to smaller active dispersers.

## Methods

### Study area

We looked for well-preserved temporary freshwater bodies from sea level up to 1,500 m a.s.l. within a similar spatial extent in two distant biogeographic regions (Neotropical and Mediterranean) with dry tropical and mediterranean climates respectively. There, we selected 30 tropical and 32 mediterranean (seasonal or semipermanent) temporary ponds in a study area of 13,000 km^2^ in Eastern Spain and 10,000 km^2^ in Northern Costa Rica. The maximum distances between ponds were 209 and 252 km in Spain and Costa Rica, respectively (Figure 1). The selected ponds encompass different typologies including peripheral areas of coastal wetlands, inland shallow lakes, interdunal slacks and small naturalized farmland ponds. All ponds are shallow (< 2 meters), fresh to oligohaline waters (< 7000 μS/cm) and vary in their hydroperiod lengths, from six months to some years. The tropical ponds experience almost constant warm temperatures during the whole year (24.9 ± 1.3 °C) and high but variable precipitation (2486 ± 934.7 mm). The mediterranean ponds are distributed in a region with warm annual mean temperature (12.9 ± 3.0 °C) and reduced annual precipitation (537 ± 68.3 mm) (Fick and Hijmans 2017). We sampled the ponds at the beginning of the hydroperiod (two weeks after infilling, during the rainy season), in May 2017 for the selected tropical ponds and January 2018 for the mediterranean ponds.

**Figure 1:**
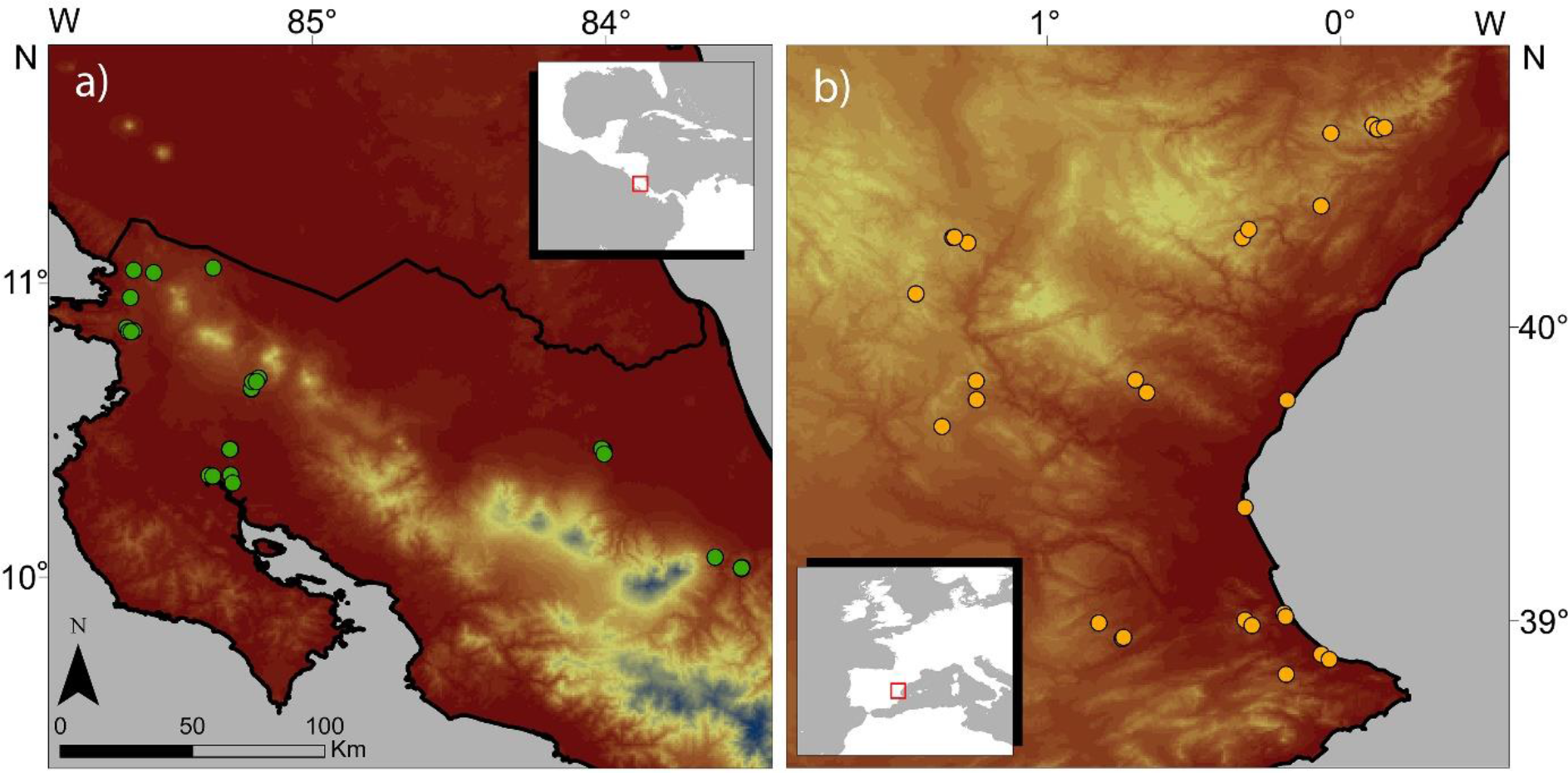
Map of the study area with the location of the ponds sampled in each region: Costa Rica (a) and eastern Spain (b).

### Environmental characterization

We measured limnological variables *in situ* including water transparency, which was measured with a Snell tube (Van de Meutter et al. 2006), as well as water temperature, pH, electrical conductivity and dissolved oxygen concentration, determined using a WTW® Multi 3430 Multiparameter Meter (with WTW® SenTix 940 for pH, TetraCon 925 for conductivity and FDO 925 for oxygen concentration). Unfiltered water samples were taken for further volumetric analyses of chloride and alkalinity, and filtered water for photometric determination of ammonium, nitrite, nitrate, sulphate and phosphate (SpectroquantMerk® and AquaMerk® test kits and T90+UV/VIS Spectrophotometer) in the lab. The water used for nutrient content analyses was filtered *in situ* through Whatmann® GF/F filters. We extracted chlorophyll-*a* from these GF/F filters with acetone 90% and analyzed the extract by spectrophotometry (Jeffrey and Humphrey 1975). Hydrogeomorphological variables included maximum and average depth, measured with a graduated stick, so as maximum and minimum diameter, surface area and shoreline development, as 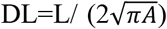, where L=perimeter and A=area (Aronow 1984), using Google Earth Pro 7.3.2.5776 (Google Inc.). We took six random sediment samples at different depths to analyze its granulometry (following De Vaasma 2010), and organic and carbonate content (according to Heiri et al. 2001); we estimated the proportion of dry weight represented by different grain sizes (>2 mm, 2-1 mm, 1-0.5 mm, 0.5-0.25 mm, 0.25-0.063 mm, 0.063-0.036 mm, <0.036 mm), and by organic matter and mineral carbonates. We estimated the hydrological regime (seasonal or semipermanent) and the main water origin (rain, streams or phreatic inputs) from information by local experts, direct observation and maps. Biotic variables (in addition to chlorophyll-*a*) included the percentage of surface area of the pond covered by submerged, floating and emergent vegetation (visually estimated), the presence of fish and the presence of livestock. Landscape metrics were estimated using Google Earth, including elevation above sea level and the percentage of different types of land cover (agricultural fields, low grass, high grass, scrub, forest or buildings) in a circular area of 100 m in diameter of the landscape closest to the sampling point. Land cover diversity was measured using the Shannon index of the different land cover types. Finally, climatic variables were extracted from WorldClim (Fick and Hijmans 2017) to obtain mean annual, maximum and minimum temperatures, temperature range, mean annual precipitation, and precipitation seasonality for each pond, using the software ArcGIS 10.3 (ESRI 2014).

### Biological communities

For each pond, we characterized prokaryotic (Bacteria and Archaea), phytoplankton, zooplankton, benthic invertebrate, amphibian and bird communities. For prokaryotic organisms, two samples were obtained: a 1-L sterilized bottle with pond water and a microtube with 2 mL of pond wet sediment. Afterwards, in the laboratory, we filtered the water through 3 μm pore polycarbonate filters (Nucleopore, Whatman) to remove larger particles. The remaining water was filtered again through 0.2 μm pore polycarbonate filters (Nucleopore, Whatman) to concentrate samples for subsequent DNA analysis. Filters and sediment samples were cold-stored in microtubes filled with RNAlater™ reagent, until further processing. DNA extractions, PCR and bioinformatics for taxonomic assignments were made following Picazo et al. (2019).

Phytoplankton samples were obtained directly from the water column at the center of the pond (100 mL stored in amber colored glass bottles) and fixed with 3 mL of Lugol’s solution. Phytoplankton taxa were identified to species level whenever possible using mainly Huber-Pestalozzi (1976-1982) and Wolowski and Hindák (2005). Zooplankton samples were taken dragging a hand net (63 μm mesh-size) through 10-20 m of water column, whenever possible, integrating all different microhabitats and depths. Samples were fixed in formaldehyde 4% (final concentration, v/v) and identified at the species level, whenever possible, following Koste (1978) and Segers (1995) for rotifers, Alonso (1996), Elías Gutiérrez et al. (2008), and references therein for branchiopods, Dussart (1967; 1969) and Elías-Gutiérrez et al. (2008) for copepods, and Blędzki and Rybak (2016) for both branchiopods and copepods.

Benthic invertebrate communities were sampled using a hand net (20 × 20 cm, 250 μm mesh-size). About 10 m in total (if ponds were large enough) were sampled across all microhabitats. Samples were fixed in ethanol 96% and invertebrates identified to the lowest taxonomic rank possible mostly following Wiederholm (1983), Tachet et al. (2010), Thorp and Covich (2010) and Springer et al. (2010), and references therein, for macroinvertebrates, and Meisch (2000), Karanovic (2012) and references therein for Ostracoda.

The occurrence of amphibian species in the ponds was registered *in situ* by noting the presence of eggs, tadpoles, adults and calls. Due to the low species richness and the easy detectability of species in the mediterranean ponds, we surveyed each pond with a hand net (800 cm^2^, 2 mm mesh pore) with a constant effort of ten minutes, and we examined the surroundings of the pond for ten more minutes. When *in situ* identification of larvae was not possible, we identified them in the laboratory by their oral disk. In tropical ponds, with higher species richness and lower detectability, we performed night surveys looking for individuals in the surroundings of each pond with an effective effort of up to 2 hours per pond, avoiding full moon nights (Wilkinson 2015). Bird surveys were performed by 15-minute point-counts per pond in which we recorded species presences identified either visually or by their calls (Ralph et al. 1995).

### Statistical analyses

We quantified the role of environmental selection, ecological drift and dispersal limitation, by partitioning metacommunity variation (Peres-Neto et al. 2006) of each taxon and region between environmental variables and spatial functions. The groups considered were Archaea from the sediment, Archaea from the water, Bacteria from the sediment, Bacteria from the water, phytoplankton, Rotifera, microcrustaceans, macroinvertebrates, Amphibia and Aves. We also analyzed subgroups of these organisms separately due to major taxonomic and trophic differences among them. Phytoplankton was therefore split as Cyanobacteria, Chlorophyceae, Bacillariophyceae (diatoms) and mixotrophic flagellate phytoplankton (including Chrysophyceae, Cryptophyta, Euglenophyta and Dinoflagellata). Microcrustaceans were split in Branchiopoda, Copepoda and Ostracoda, and macroinvertebrates in Mollusca and Insecta. Moreover, insects were divided in Palaeoptera (including Odonata and Ephemeroptera), Heteroptera, Coleoptera and Diptera (considering also Chironomidae separately as another subgroup). Separately, groups and subgroups accounted for 24 species matrices based on presence-absence data for each region.

For each group (and subgroup) and region, we performed variation partitioning via Generalized Additive Models (GAMs; Wood 2011), recently used to explore metacommunity variation (Viana et al. 2021). Despite its popularity in community analyses, the performance of Redundancy Analyses (RDA) is weak when species respond non-linearly to environmental variation (e.g., Makarenkov and Legendre 2002).To tackle its limitations, we used GAMs as a non-linear modeling framework considering both environmental and spatial variables, allowing a better fit to the data.

Response matrices in GAMs were composed of predicted values of the presence/absence data, obtained from Generalized Linear Latent Variable Models (GLLVM with binomial distribution) with the lowest AIC (Niku et al. 2017). This allowed us to concentrate on the common sources of variation among species and reduce computational time from extracting GAMs for a really large number of taxa. Prior to these analyses, environmental variables were inspected and transformed either logarithmically or with the arcsine of the square root, depending on the distribution of the raw data, in order to reduce skewness and leverage of extreme values (McDonald 2014). When applying GAMs, due to the small sampling size (30 tropical and 32 mediterranean ponds), we were limited to a maximum number of three variables, to allow GAMs generate nine splines per variable (3 variables × 9 splines = 27), maximizing the response of the selected variables. As a consequence, before executing GAMs (using a quasi-binomial distribution), we reduced the number of environmental predictors by using the first three principal components (PCs) in a PCA of all variables. As spatial predictors, we used spatial functions (the default splines in the mgcv R package) calculated from longitude and latitude coordinates.

We reduced the number of environmental and spatial predictors introduced in GAMs by means of forward selection, with a double-stopping criterion (Blanchet et al. 2008). First, we selected only significant variables (p-value<0.05). Second, the adjusted R^2^ of the selected variables had to be lower than the adjusted R^2^ of the model with the whole set of variables. This procedure was repeated for the environmental variables (three PCs) and spatial variables (latitude and longitude) separately. We included at least one environmental or spatial variable in rare cases that the double-stopping criterion was not satisfied, as long as the selected variable was significant. If more than three variables were selected, we manually reduced the number of variables (either spatial or environmental) by removing the selected variables which less contributed to explaining the observed variation, until reaching the maximum number of three selected variables. As a result, GAMs provide information on the total proportion of metacommunity variation explained by environmental (E) and spatial (S) factors together (E+S) and separately, estimating the pure environmental fraction (E|S; associated with environmental selection), the pure spatial fraction (S|E; that can be interpreted as variation due to dispersal processes assuming that environmental variation was mostly accounted for) and the overlap of environment and space (E∩S; which represents environmental variation that is spatially structured). The unexplained proportion or residuals represent common and uncommon sources of variation not explained by (measured) environmental and spatial variation.

The contribution (R^2^) of environmental variation to species distributions (i.e., species sorting via environmental selection) can be inflated when residuals and response variables (i.e., species distributions here) are both autocorrelated. Traditionally, ecologists remove the spatialized component of the environment via variation partitioning, focusing then solely on the E|S fraction. This procedure, however, may reduce estimates of the importance of environmental selection because even the spatialized component of the environment that may not bias estimates of variation partitioning is eliminated from the E|S fraction (see Clappe et al. 2018 for a discussion). To assess the importance of environmental drivers (spatialized and non-spatialized) we used the correction method for the environmental component as described in Clappe et al. (2018). This method produces unbiased estimates of the environmental contribution even under spatial autocorrelation of residuals. In order to make results comparable among taxa and region, we transformed the fractions of explained variation into the relative proportion of purely environmental effects (E|S/(E+S)), purely spatial effects (S|E/(E+S)) and spatialized environment (E∩S/(E+S)). Finally, we carried out a test of homogeneity of multivariate dispersion (PERMDISP; Anderson 2006) to contrast the environmental heterogeneity of both regions (tropical *versus* mediterranean) using the whole set of measured environmental variables, as well as a matrix of just limnological variables (which are expected to be more local) and a matrix of just climatic variables (which are expected to be more spatialized). All analyses were performed with R (v4.0.2; R Core Team 2021) and R packages *vegan* (Oksanen et al. 2019), *ade4* (Bougeard and Dray 2018), *adespatial* (Dray et al. 2021), *gllvm* (Niku et al. 2020) and *mgcv* (Wood 2017). Environmental characterization and community matrices can be found in Figshare (doi.org/10.6084/m9.figshare.14644608.v4). Codes for variation partitioning analyses can be found here: github.com/Angel-Galvez/Bacteria-to-birds.

## Results

The total proportion of explained variation by environmental and spatial components (E+S) ranged from 0.10 for mollusks to 0.77 for chironomids in the tropical ponds (average 0.42 ± 0.18). In the mediterranean ponds, the explained variation varied from 0.08 for sediment Archaea to 0.93 for birds (average 0.54 ± 0.23). Results of these variation partitioning analyses can be found in Appendix S1: Table S1 and Figure S1. In addition, results of PERMDISP showed non-significant differences in environmental heterogeneity between regions for the whole environmental dataset (p-value=0.303, 6.159 average distance to centroid in mediterranean ponds, 5.613 in tropical ponds). However, mediterranean ponds showed significantly higher limnological heterogeneity (p-value=0.002, 3.113 average distance to centroid in mediterranean ponds, 1.752 in tropical ponds), while tropical ponds exhibited significantly higher climatic heterogeneity (p-value=0.005, 1.311 average distance to centroid in tropical ponds, 0.921 in mediterranean ponds).

When we compared the relative contribution of environmental and spatial components for each group between biogeographic regions (Figure 2), we clearly found higher pure environmental effects (E|S) in mediterranean than in tropical metacommunities. In contrast, the pure spatial fraction (S|E) was generally higher in the tropical metacommunities, despite some mediterranean groups exhibited an important spatial control (Ostracoda, Heteroptera, Palaeoptera, diatoms and the mixotrophic flagellate phytoplankton). The overlap of environment and space (E∩S) was greater in tropical ponds. The total environmental fraction (E) was higher in the mediterranean-climate region, while total spatial fraction was higher in the tropical region. Pure environment (E|S) explained a higher proportion of variance than purely spatial component (S|E) in mediterranean ponds, while pure spatial and purely environmental effects were similar in tropical ponds.

**Figure 2:**
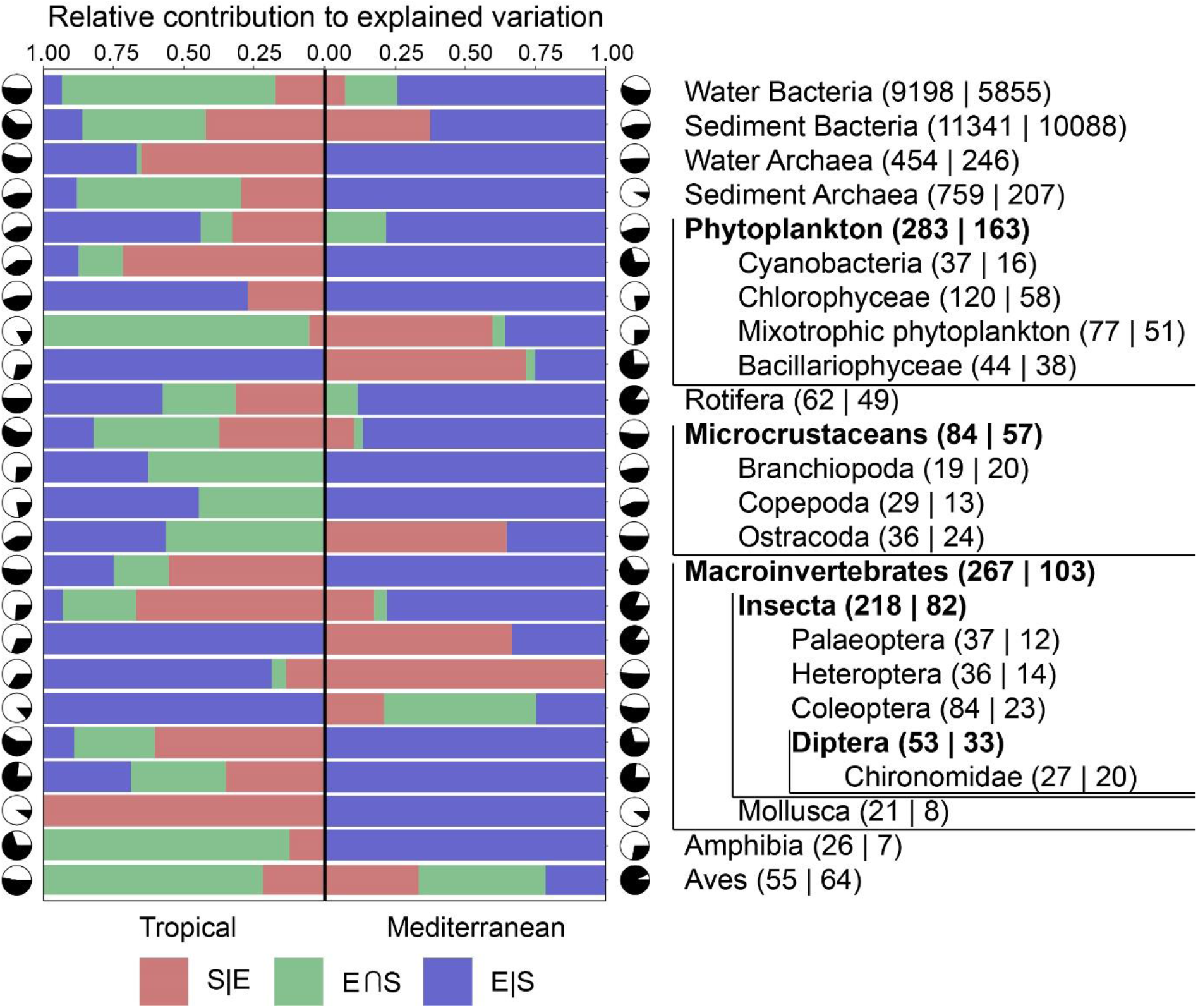
Results of variation partitioning analysis for each group of organisms in tropical and mediterranean metacommunities. The relative contribution of pure environment (E|S/(E+S)), spatialized environment (E∩S/(E+S)) and space (S|E/(E+S)) are represented with a different color. Black portions in pie charts represent the total proportions of explained variation, and white portions the residual variation. Groups in bold type include species from the groups enclosed in the corresponding following indented line(s). Number of identified taxonomic units, i.e. species in most groups, are shown between brackets (tropical | mediterranean) next to each group label.

Regarding the relative role of space and environment within organisms and across regions (Figure 3), the pure environmental fraction generally plays a greater role in mediterranean groups than in their tropical counterparts. Pure spatial effects are important structuring metacommunities of even small organisms such as Bacteria and Archaea. As for the spatialized environment and the pure spatial fraction, we found tropical groups to be more influenced by the environmental and spatial overlap or by purely spatial effects than their mediterranean homologous groups. However, the relative contributions of these components do not follow any trend on body (or propagule) size neither for active nor passive dispersers. If we compare the relative contributions of space and environment on groups of passive and active dispersers (Figure 4), we find similar fractions explained by purely environmental, spatialized environmental and spatial effects in both passive and active tropical dispersers. Despite large differences in environmental and spatial contributions in the mediterranean ponds, both passive and active dispersers behave similarly, although Mediterranean active dispersers are more affected by pure environmental factors, and more distant in that matter from the tropical groups, than passive ones. Thus, passive and active dispersers mostly show similar relative contributions of environmental and spatial effects within regions, but differently across regions; environmental effects are higher in mediterranean than in tropical ponds, while the spatial component is slightly higher in the tropical than in the mediterranean-climate region.

**Figure 3:**
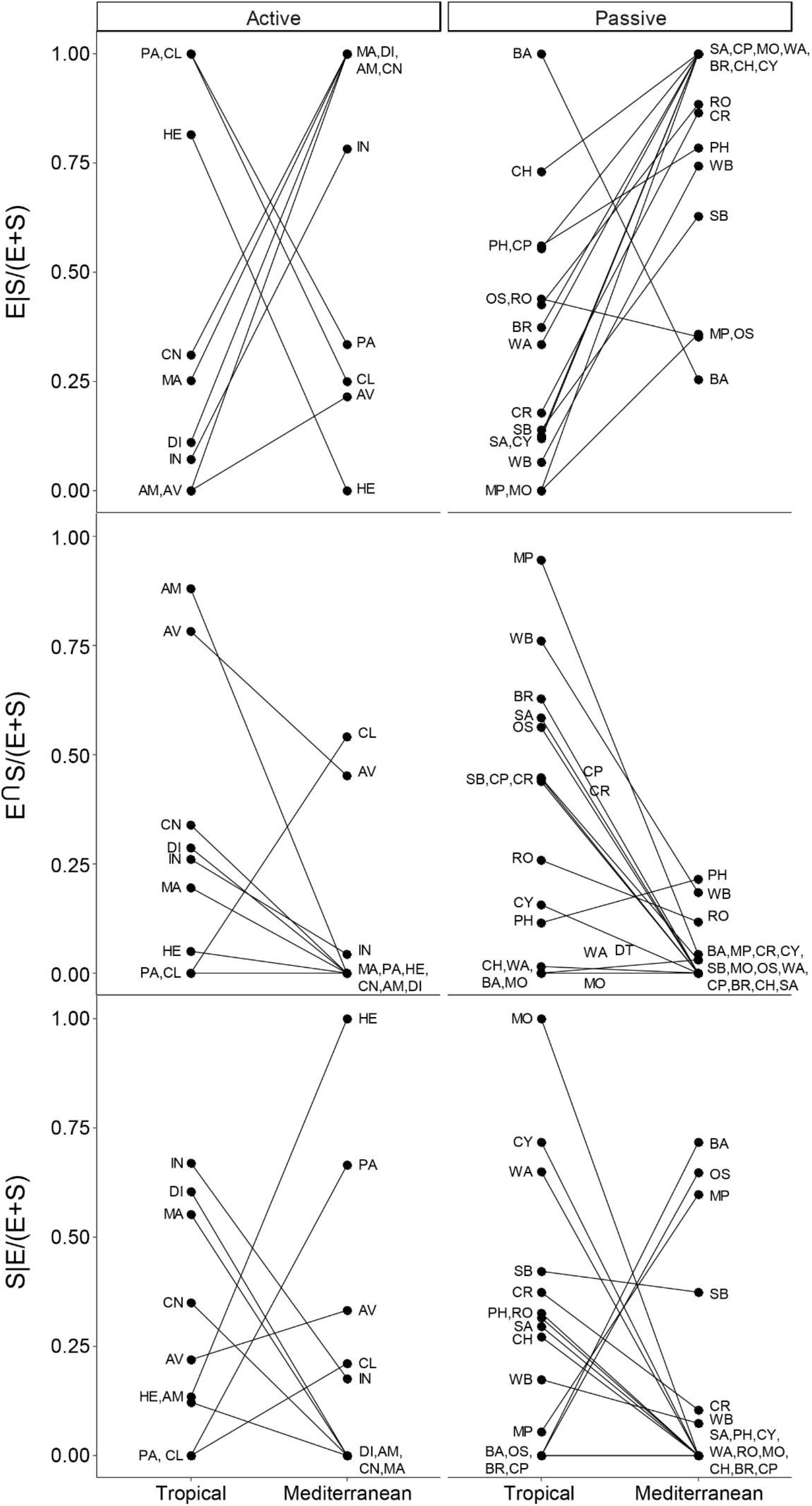
Relative contributions to total explained variation by the pure environmental component (E|S/(E+S), spatialized environment (E∩S/(E+S)) and pure spatial effects (S|E/(E+S)), for each group of organisms, and according to geographic region and dispersal mode. WB: water Bacteria, SB: sediment Bacteria, WA: water Archaea, SA: sediment Archaea, PH: phytoplankton, CY: Cyanobacteria, CH: Chlorophyceae, MP: mixotrophic flagellate phytoplankton, BA: Bacillariophyceae, RO: Rotifera, CR: microcrustaceans, BR: Branchiopoda, CP: Copepoda, OS: Ostracoda, MA: macroinvertebrates, IN: Insecta, PA: Palaeoptera, HE: Heteroptera, CL: Coleoptera, DI: Diptera, CN: Chironomidae, MO: Mollusca, AM: Amphibia, AV: Aves. Black lines connect the same groups from different regions.

**Figure 4:**
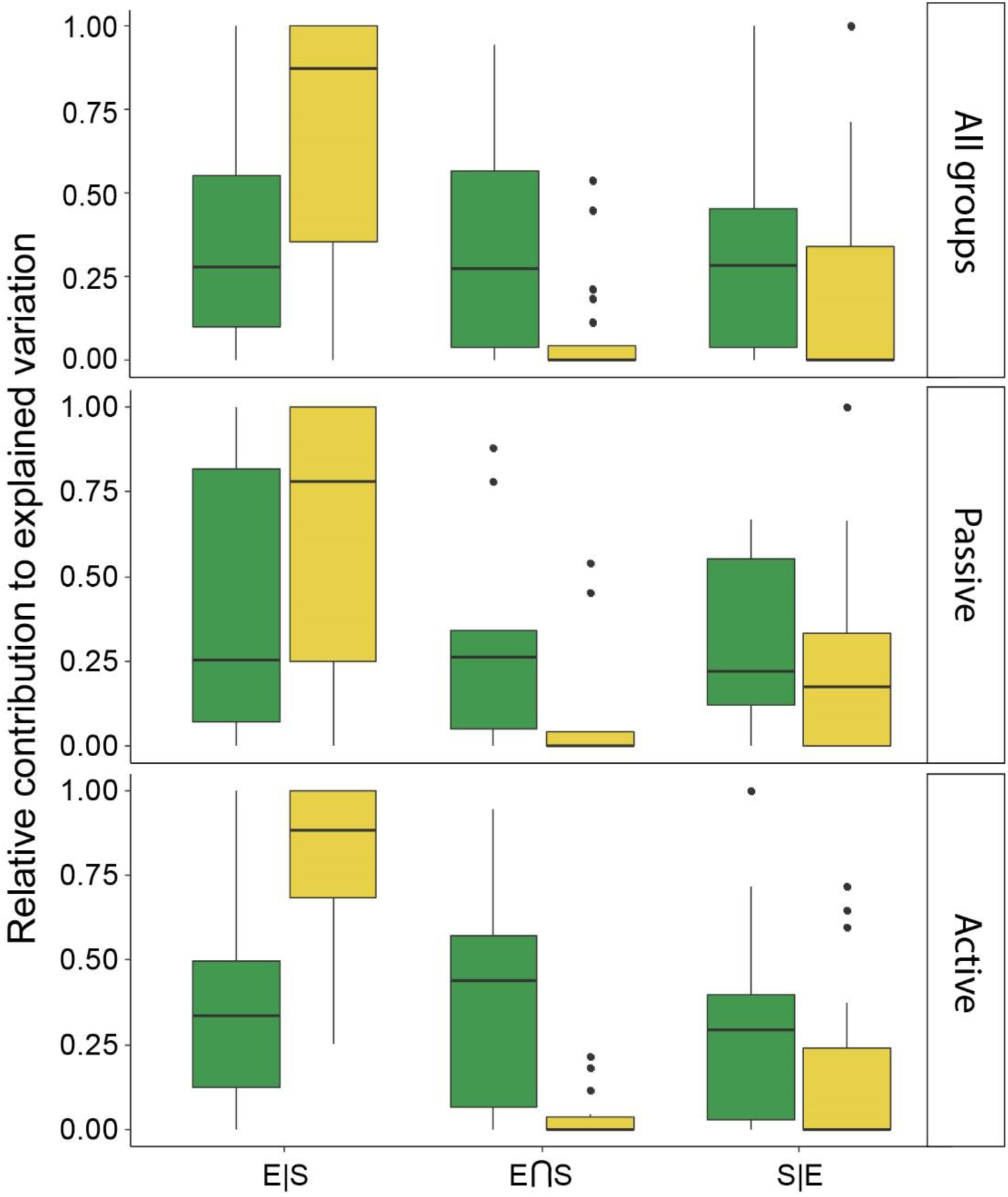
Relative contribution to total explained variation of the pure environmental component (E|S/(E+S)), the spatialized environmental component (E∩S/(E+S)), and the pure spatial component (S|E/(E+S)) for all groups or separately for passively and actively dispersing organisms. Green and yellow colors represent tropical and mediterranean ponds, respectively.

## Discussion

In this study, we set out to contrast the environmental and spatial contributions on metacommunity structure among a wide range of taxa inhabiting temporary ponds in two distant biogeographic regions, Neotropical and Mediterranean, with large differences in climate and biodiversity. Overall, we found a lower proportion of explained variation in tropical than in mediterranean pond metacommunities, considering both environmental and spatial effects across the diverse array of studied organisms. This is consistent with the expectation that more productive environments in the tropics, with greater and constant high temperatures, might display more stochasticity in community assembly than less productive ones and with higher temperature fluctuations, such as ponds in mediterranean areas, where metacommunities might be modulated more tightly by environmental determinism (Chase 2010).

Our results show that both environmental selection and dispersal processes (here interpreted as spatial variation) played an important role in the assembly of pond metacommunities in both regions. However, pure environmental effects were stronger in mediterranean compared to tropical pond metacommunities, as previously found for vegetation and bacterioplankton metacommunities (Myers et al. 2013; Souffreau et al. 2015). Given the higher limnological heterogeneity across the studied mediterranean water bodies, these results were therefore not unexpected (Ai et al. 2013), as larger gradients can generate more environmental space for divergently specialized organisms. So, even if there are more species in the tropical ponds, given their shorter environmental gradient and reduced species sorting, we expect to find a higher degree of ecological redundancy and stochasticity in these ponds. However, we cannot discard unmeasured important environmental factors or intense biotic interactions, which could play important roles in tropical metacommunities (Roslin et al. 2017; Leibold and Chase 2018).

We hypothesized spatial effects to be lower in the tropical metacommunity, due to a shift in connectivity during the rainy season, which should lead to community homogenization (Thomaz et al. 2007; Rojo et al. 2016; Brasil et al. 2020), in comparison to the reduced precipitation in the mediterranean region, which is expected to make ponds more isolated one from another. Unexpectedly, we found the opposite pattern. Low spatial effects in mediterranean ponds might reflect historical connectivity due to transhumance (Incagnone et al. 2014) or intense movements of potential vectors including birds and mammals among ponds (Frisch et al. 2007; Vanschoenwinkel et al. 2008; Valls et al. 2016, 2017). The strong spatial effects in the tropical metacommunity might also be partly explained by the sampling period, given that at early stages of the hydroperiod, connectivity is not yet at maximum. It is possible that during early stages of the hydroperiod, dispersal constraints might be equally strong for aquatic organisms in both regions. Additionally, annual floods during the wet season drive connectivity changes, homogenizing both environmental conditions and biological communities (Thomaz et al. 2007; Brasil et al. 2020). Thus, early-hydroperiod spatial patterns in small tropical organisms can be the result of historical contingencies (Castillo-Escrivà et al. 2017b), such as sequential flooding events (or heavy rain periods) that shifted connectivity at regional scale. These events may increase the spatial structure of passively dispersed organisms in more connected areas compared to other further away and more isolated regions.

In addition to hydrology, orography could be a stronger dispersal barrier in the tropics than in temperate zones. Dispersal of tropical species, adapted to homogeneous temperature regimes, could be more limited by sharp altitudinal climate shifts than temperate species, adapted to a wider range of temperatures (Janzen 1967). The tropical ponds in our study are distributed across two watersheds separated by the Continental Divide, with wide differences in rainfall amount and seasonality, which can have varying impacts on the distributions of some species (Chandler and King 2011). This orographic barrier and the associated abrupt climatic transition are probably characterized by the shared components between environmental and spatial components (E∩S), which was found to be higher for the tropical ponds, where climatic heterogeneity was also higher. Thus, reduced limnological gradients and orographic barriers can lead to dispersal limitation (Janzen 1967) at larger scales, and higher regional connectivity can lead to dispersal surpluses (Thomaz et al. 2007; Heino et al. 2015; Brasil et al. 2020) at smaller scales in the tropical area. These could explain the proportionally greater role of space (assuming they represent dispersal signatures) in tropical than in mediterranean metacommunities.

Some studies have found that the degree of variation of spatial *versus* environmental effects on metacommunity structure strongly depends on propagule size and the dispersal mode of organisms (De Bie et al. 2012; Heino et al. 2015). However, we found it to be very variable for the same group between geographical settings (i.e., mediterranean *versus* tropical). A significant spatial effect was found even in the smallest studied organisms, including prokaryotes, planktonic protists and small metazoans. Organisms smaller than 1 mm have traditionally been considered ubiquitous, lacking biogeographical effects (Finlay 2002), although more recent analyses suggest there is no strong support for or against this statement (Fenchel et al. 2019). In this sense, we found significant spatial effects in both active and small passive dispersers, including Bacteria and Archaea, supporting the view that microorganisms are subjected to basically the same processes as those affecting macroorganims (Hortal 2011). For instance, the effects of extreme isolation unveil dispersal limitation allowing allopatric differentiation even in bacterial subspecies, as it was shown for Maritime Antarctic lakes (Hahn et al, 2015). Indeed, we did not find the expected differences in spatial and environmental effects between different body-sized, active or passive dispersers from the same region (De Bie et al. 2012; Padial et al. 2014), as neither did Heino et al. (2012) nor Schulz et al. (2012). Passive and active dispersers within the same region barely differed between them in their spatial and environmental effects, suggesting that spatial and environmental constraints are similar for organisms with different dispersal abilities (Schulz et al. 2012). Large fractions of unexplained community variation remain, however, suggesting an important role for the elements of chance or randomness (Jeffries 1989). Stochastic processes and priority effects are expected to be strong at the beginning of the hydroperiod due to egg-bank hatching, before abiotic and biotic sorting of species can effectively act as filters in structuring local communities (Antón-Pardo et al. 2016; Castillo-Escrivà et al. 2017c; Mahaut et al. 2018). This results in increased unexplained variation, with model residuals accounting for a higher proportion in the tropical metacommunity, probably related to its higher species richness, but also because of lower limnological heterogeneity (Leibold and Chase 2018), as shown by the homogeneity test of multivariate dispersion. Higher limnological heterogeneity may therefore account for a stronger influence of environmental selection processes in the mediterranean ponds and we may also expect a higher influence of ecological redundancy in the richer tropical communities, under shorter environmental gradients. In addition, biotic interactions are supposed to be stronger in the tropics (Roslin et al. 2017). Furthermore, unmeasured biotic variables, which have been neglected for years in metacommunity studies, together with the loss of environmental information due to model selection in GAMs, can give rise to misleadingly high proportions of unexplained residual variation.

Despite some potential limitations, we found that niche and dispersal-related processes played an important role in structuring metacommunities of tropical and temperate temporary ponds, although environmental effects dominated in the mediterranean region (i.e. the more heterogeneous limnological environment). Contrary to expectations underlying dispersal processes, the type of dispersal ability of the studied taxa (active or passive) did not relate to the proportion of spatial and environmental control on their metacommunities. We found no general pattern among taxonomic groups between regions, with very variable and idiosyncratic responses. Organisms are generally more environmentally than spatially controlled, and more so in the temperate zone than in the tropics. This is however a snapshot study at a time characterized by high stochasticity at the beginning of the hydroperiod. In order to further disentangle the actual role of each process shaping metacommunities, we need temporal tracking of these metacommunities to better quantify not only the relative role of environmental and spatial components along a temporal series, but also of time *per se* in structuring metacommunities in highly dynamic ecosystems.

## Acknowledgments

This study was sustained by the Spanish Ministry of Economy, Industry and Competitivity–AEI, and FEDER (EU), through project METACOM-SET (CGL2016-78260-P). Ángel Gálvez was also supported by an FPI fellowship BES-2017-080022 from the Spanish Ministry of Economy, Industry and Competitiveness. Berenice de Manuel is thanked for her help in laboratory analyses.

## Appendix S1 Metacommunities from bacteria to birds: stronger environmental selection in mediterranean than in tropical ponds

**Table S1:**
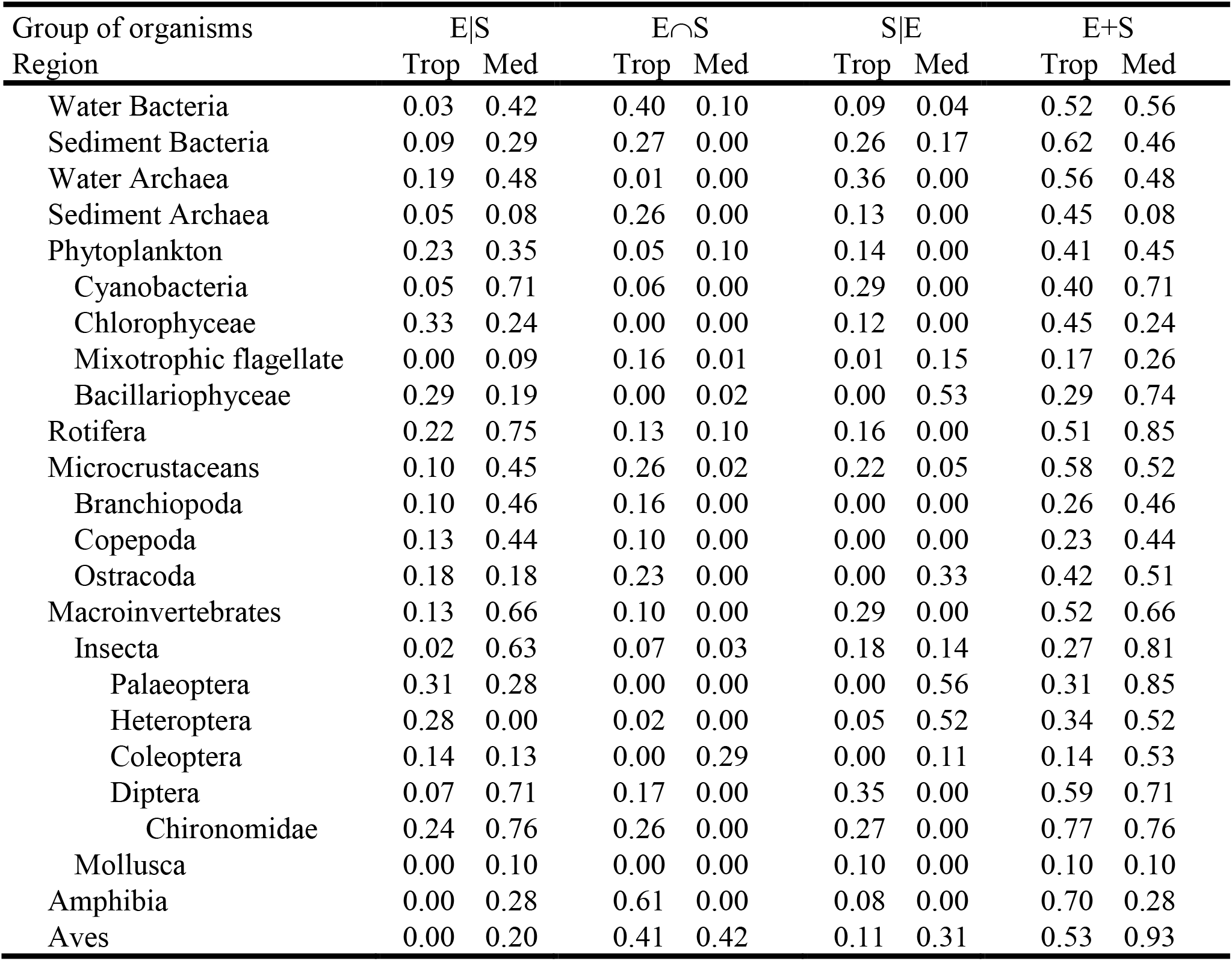
Results of variation partitioning analysis for each group. The table shows the proportion of variation (R^2^_adj_) explained by each pure component (E|S and S|E), the overlaps between components or spatialized environment (E∩S), and the total explained variation (E+S) in tropical and mediterranean metacommunities.

**Figure S1:**
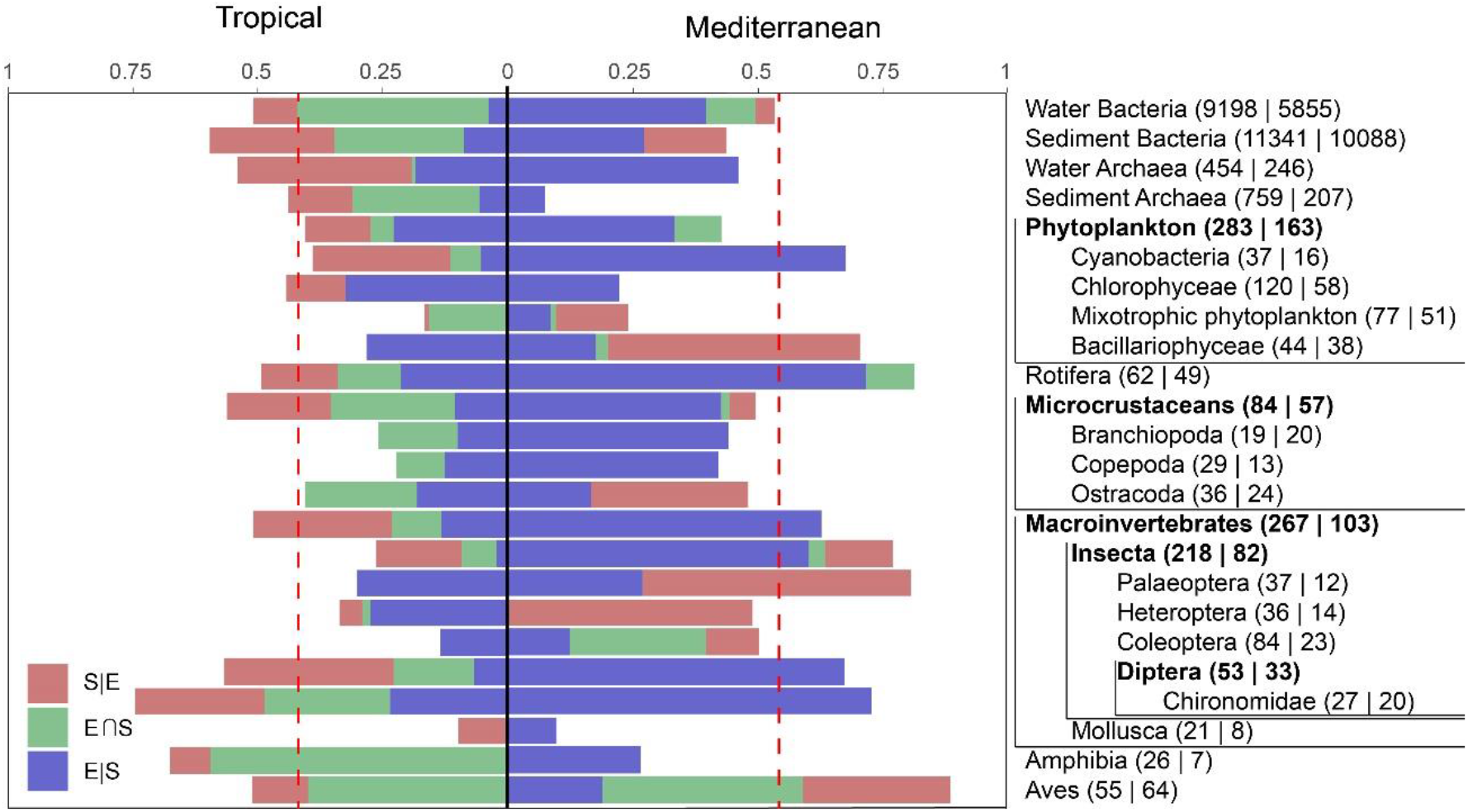
Results of variation partitioning analysis for each group of organisms in tropical and Mediterranean metacommunities. The proportion of variation explained by each component is represented with a different color. Taxa in bold type include species from the following groups enclosed in the corresponding line. Red dashed line represents average total explained variation. Number of identified species (Tropical | Mediterranean species) are shown next to each group label.

